# Systematic optimization of *Caenorhabditis elegans* cryopreservation

**DOI:** 10.1101/2025.07.19.665704

**Authors:** Sajal Agrawal, Ashutosh Karharia, Kamesh R. Babu

## Abstract

*Caenorhabditis elegans (C. elegans)* is a non-parasitic roundworm widely utilized as a versatile model organism for studying fundamental biological processes. Despite the availability of multiple cryopreservation methods, variations in the selection of developmental stage, cryoprotectant composition, and storage conditions may sometimes cause inconsistencies and uncertainty among researchers. In this study, we followed a step-by-step approach to optimize the *C. elegans* cryopreservation process. Firstly, different larval stages (L1-gravid adult) of *C. elegans* were cryopreserved using a fixed concentration of trehalose-DMSO and glycerol, followed by the post-thaw survival rate assessment. We found that the starved L1 larvae has the highest survival rate among other developmental stages. Subsequently, the starved L1 larvae was cryopreserved using different concentrations (5%, 10%, and 15%) of DMSO and glycerol and found that 5% DMSO has the most effective cryoprotection. Finally, the impact of storage conditions on worm viability was evaluated by placing cryogenic vials in different storage boxes, including standard cardboard cryogenic boxes, Styrofoam boxes, and isopropanol-based freezing containers, and found that standard cardboard cryogenic boxes provided higher worm viability. The finalized protocol was then validated across multiple mutant and transgenic strains to confirm its reproducibility and broad suitability. Taken together, our results demonstrate that starved L1 larvae cryopreserved with 5% DMSO using a standard cardboard cryogenic box provided the highest survival rate (>90%) across wild-type, mutant, and transgenic strains. Here we propose a simple, reproducible, and strain-independent protocol for cryopreservation of *C. elegans*. The proposed protocol can be easily adopted in *C. elegans* research laboratories and stock centers globally.

## 1. Introduction

*Caenorhabditis elegans* (*C. elegans*) is a free-living, non-parasitic nematode widely recognized as a powerful model organism in biomedical and genetic research due to its well-characterized genome, short life cycle, and ease of maintenance^1,2^. It has been extensively used to investigate developmental biology^3^, neurobiology^4^, aging^5^, and disease mechanisms^6-9^. Given the growing use of *C. elegans* in functional genomics^10,11^, mutant screening^12,13^, and transgenic studies^14,15^, reliable long-term preservation of various strains has become essential for laboratory workflows and stock center operations.

Cryopreservation offers an efficient solution for long-term storage of *C. elegans* strains. The earliest standardized method for cryopreserving *C. elegans* was introduced by Sydney Brenner in 1974 as part of his seminal work establishing the nematode as a model organism. Brenner’s protocol involved the use of a glycerol-based freezing solution, where worms of mixed larval stages were suspended in S buffer supplemented with 15% glycerol and slowly frozen (-1 °C/min) at -80□°C in cryogenic vials and later placed in a liquid nitrogen container^16^. The protocol is widely regarded as the standard for *C. elegans* freezing and is officially recommended by *WormBook*, a comprehensive online review of *C. elegans* biology^17,18^. Many laboratories across the world continue to use this as a standard protocol for *C. elegans* cryopreservation in their research workflows^19-22^. Ironically, this approach enabled long-term storage of worm stocks but often resulted in variable survival rates depending on worm stage^16^. Scientists have also explored the use of alternative cryoprotective agents for the nematode cryopreservation. One of the earliest systematic studies was conducted in 1975 by Haight *et al*., who evaluated the freeze-thaw survival of *Caenorhabditis briggsae*, a species closely related to *C. elegans*, using dimethyl sulfoxide (DMSO) as a cryoprotectant^23^. Moreover, a subsequent study has demonstrated that DMSO offered superior preservation of *C. elegans* viability when compared to glycerol^24^. Later, K. O’Connell proposed a combination of trehalose and DMSO as an alternative cryoprotectant, which was claimed more effective than the conventional glycerol-based freezing solution^25^. In contrast to K. O’Connell’s claim, a study by McClanahan *et al*., reported that glycerol-based cryoprotectant was equally effective as trehalose-DMSO for *C. elegans* cryopreservation^22^.

Studies suggest that the *C. elegans* post-thaw survival rates depends on their nutritional status and developmental stage. Cryopreservation using Brenner’s glycerol-based protocol has been reported to be more effective in freshly starved early larval stages (L1-L2) compared to well fed worms or adults^16,17^. Similarly, Haight *et al*., reported that L2 juveniles of *Caenorhabditis briggsae* survived the best among other developmental stages (Egg, L3, L4, and adult) when cryopreserved using DMSO^23^. Furthermore, Hayashi *et al*., demonstrated that the L1 stage exhibited the highest post-thaw survival when cryopreserved with either DMSO or glycerol, compared to later larval stages (L2-L4) of *C. elegans*^24^. In contrast to these findings, McClanahan *et al*., observed that the post-thaw survival of early larval stages (L1-L3) was comparable to that of a mixed-stage population when cryopreserved using both glycerol-based and trehalose-DMSO solutions^22^. Moreover, studies also suggest that the freezing rate affects the worm’s viability during cryopreservation^23,24^, and recommends slow freezing at the rate of -0.2 °C/min^23^ or -1 °C/min^16,17^ to achieve better post-thaw survival. To achieve the slow freezing rate during cryopreservation researchers typically stores the cryogenic vials either in Styrofoam boxes^17,21,22,24^ or isopropanol-based freezing containers, which are engineered to achieve a uniform -1 °C/min cooling rate^25^.

Altogether, these studies suggest that the efficacy of *C. elegans* cryopreservation is influenced by multiple factors including developmental stage, cryoprotectant composition and storage conditions. Existing protocols vary in their choice of developmental stage, cryoprotectant composition, and storage conditions, often may lead to inconsistencies in post-thaw viability and reproducibility. Most commonly, laboratories rely on freezing mixed-stage populations using conventional glycerol-based freezing solution, which may not yield consistent recovery, particularly for sensitive or transgenic strains. For example, in our laboratory, we were unable to recover the transgenic mutant strain SJ4005 after cryopreservation using the standard glycerol-based freezing protocol, resulting in complete loss of the line. Such experiences, along with contrasting findings across studies, emphasize the need for further comparative studies to establish standardized protocol for the *C. elegans* cryopreservation. Furthermore, limited studies have validated optimized protocols across mutant and transgenic strains, which is critical for ensuring broad applicability and reproducibility.

In this study, we employed a stepwise approach to systematically optimize the cryopreservation protocol for *C. elegans*. We first evaluated the survival of different developmental stages under fixed cryoprotectant conditions, followed by optimization of DMSO and glycerol concentrations. We then assessed the effect of different storage containers on worm viability. Finally, the optimized protocol was validated across multiple mutant and transgenic strains to confirm its reproducibility and broad suitability. Our findings provide a simple, reproducible, and genotype-independent protocol for the cryopreservation of *C. elegans*, suitable for adoption by research laboratories and stock centers worldwide.

## 2. Materials and Methods

### 2.1. Reagents

#### 2.1.1. Bleaching Solution (2X)

The bleaching solution was prepared as previously described^26^. A total of 1 mL of 2X bleaching solution was freshly prepared by mixing 0.3 mL of 4% NaOCl, 0.625 mL of 1 M NaOH, and 0.125 mL of double-distilled water.

#### 2.1.2. M9 buffer

M9 buffer was prepared as previously described^26^. A total of 100 mL of M9 buffer was prepared by dissolving 0.6 g of Na_2_HPO_4_, 0.3 g of KH_2_PO_4_, and 0.5 g of NaCl in double-distilled water. Then, 0.1 mL of 1M MgSO_4_ was added, and the volume was adjusted to 100 mL with double-distilled water. The solution was autoclaved and allowed to cool to room temperature before use.

#### 2.1.3. S-basal buffer

S-basal buffer was prepared as previously described^17^. A total of 100 mL of S-basal buffer was prepared by dissolving 0.59 g of NaCl in double-distilled water. Then, 5 mL of potassium phosphate buffer (pH 6.0) was added, and the final volume was adjusted to 100 mL with double-distilled water. The solution was autoclaved and allowed to cool to room temperature before adding 0.1 mL of 5 mg/mL cholesterol in ethanol.

#### 2.1.4. S-complete buffer

S-complete buffer was prepared as previously described^17^. A total of 100 mL of S-complete buffer was prepared by mixing 1 mL of 1 M potassium citrate buffer (pH 6.0), 0.1 mL of 10X trace metal solution, 0.3 mL of 1 M CaCl_2_, and 0.3 mL of 1 M MgSO_4_ with 98.3 mL of S-basal buffer.

#### 2.1.5. Freezing solution

Freezing solution was prepared as previously described^17^. A total of 100 mL of freezing solution was prepared by mixing 15 mL of glycerol with 85 mL of S buffer (11 mL of 0.05M K_2_HPO_4_, 74 mL of 0.05M KH_2_PO_4_, and 0.5 g of NaCl).

#### 2.1.6. Trehalose-DMSO solution

The trehalose-DMSO cryoprotectant solution was prepared as previously described^25^. Briefly, 3.02□g of trehalose was dissolved in 96.46□mL of M9 buffer, followed by the addition of 3.54□mL of DMSO to make a final volume of 100□mL. The solution was then filter-sterilized using a 0.22□µm syringe filter and stored at room temperature until use.

#### 2.1.7. Preparation of DMSO and glycerol solutions at different concentrations

Cryoprotective solutions were prepared by mixing DMSO or glycerol with M9 buffer to achieve final concentrations of 5%, 10%, and 15% (v/v). Each solution was freshly prepared prior to use.

### 2.2. C. elegans strains, PFA-killed OP50 and culture conditions

All the *C. elegans* strains including N2 Bristol (wild-type), AM140, AU133, DC19, GRU101, MQ1766 and MT8735 were maintained on 60 mm nematode growth medium (NGM) plates seeded with 30 µL of PFA-killed *E. coli* OP50. To prepare PFA-killed OP50, a 500 mL culture of *E. coli* OP50 was incubated overnight (∼16 hrs) at 37°C in an orbital shaker at 200 rpm. The culture was then treated with 1.25% of paraformaldehyde (PFA) for 2 hours, followed by four washes with autoclaved double-distilled water to remove residual PFA. The OP50 cell pellet was resuspended in sterile S-complete buffer at a final concentration of 250 mg/mL (∼5×10^10^ OP50/mL). Worm cultures were grown and maintained at 20°C. Some strains were provided by the CGC, which is funded by NIH Office of Research Infrastructure Programs (P40 OD010440).

### 2.3. Synchronization of C. elegans population

Confluent 60 mm NGM plates with adult worms were washed with 1 mL of M9 buffer to collect gravid adults and laid eggs. The M9 buffer containing worms and eggs was transferred to a 15 mL Falcon tube, followed by the addition of 13 mL of M9 buffer. The suspension was centrifuged at 1500 rpm for 2 minutes at room temperature. The supernatant was carefully discarded without disturbing the worm pellet, and the washing step was repeated with 14 mL of M9 buffer until the buffer appeared clear of bacteria. To the pellet, 1 mL of M9 buffer and 1 mL of bleaching solution were added, and the mixture was vortexed at 2500 rpm for 6 minutes continuously. The reaction was stopped by adding 12 mL of M9 buffer, followed by centrifugation at 2000 rpm for 1 minute at room temperature. The pellet was washed three additional times with 14 mL of M9 buffer to remove debris. The egg pellet was resuspended in either 1 ml of M9 buffer and incubated at 20°C for 15 hours at 30 rpm.

### 2.4. Cryopreservation of C. elegans

Age-synchronized *C. elegans* worms were counted using a stereomicroscope (Nikon), and a minimum of 100 worms were transferred into cryogenic vials (Tarsons) containing the respective cryoprotectant solution. The vials were then placed into one of the following storage containers: a standard cardboard cryogenic box (Tarsons), a Styrofoam box with walls of ∼1 inch thick (as per the *Wormbook* recommendation)^17^, or an isopropanol-based freezing container (Tarsons). Subsequently, all samples were stored at –80□°C in a deep freezer (Thermo Fisher).

### 2.5. Thawing of cryogenic vials and scoring of survival rate

After one week of cryopreservation, the cryogenic vials were thawed at room temperature. The thawed worms were washed with M9 buffer to remove the cryoprotectant and transferred onto 35 mm NGM plates seeded with PFA-killed *E. coli* OP50. The plates were then incubated at 20□°C for 24 hours to allow recovery. Following incubation, worms were harvested from the plates, washed with M9 buffer, and pelleted by centrifugation at 1500 rpm for 2 mins. The worm pellet was resuspended in 250□µL of M9 buffer and placed onto a clean glass slide for survival scoring. The number of live and dead worms was assessed under a stereomicroscope (Nikon).

### 2.6. Statistical analysis

Data are presented as the mean□±□standard error of the mean (SEM) from a minimum of three independent experiments. Statistical analyses were conducted using GraphPad Prism v.9. Differences in survival curves were evaluated using two-way ANOVA, while comparisons of survival rates between starved L1 larvae preserved in 5% DMSO versus trehalose-DMSO, as well as across different storage containers were assessed using a two-tailed Student’s *t*-test. A *P*-value of < 0.05 was considered statistically significant. Each experimental replicate included at least 100 worms. Statistical significance indicators are described in the respective figure legends.

## 3. Results

### 3.1. Starved L1 larvae survive better than other developmental stages of C. elegans during cryopreservation

To determine which developmental stage of *C. elegans* survives best during cryopreservation, we cryopreserved age-synchronized starved L1 larvae, and fed larvae including L1, L2, L3, L4, young adult, and gravid adult stages of the wild-type N2 strain using the well-known standard freezing solution and trehalose-DMSO solution. The cryogenic vials containing worms in the respective cryoprotectant solutions were placed in a standard cardboard cryogenic box and stored at -80□°C. After one week of cryopreservation, the vials were thawed at room temperature, and worm survival was assessed. We observed that starved L1 larvae exhibited the highest survival rate compared to all other developmental stages in both trehalose-DMSO (∼86%) and freezing solution (∼63%) conditions (Fig. 1A). Moreover, worms cryopreserved with trehalose-DMSO solution showed significantly higher survival rates across all developmental stages than those preserved with the freezing solution *(F(1, 112) = 498*.*9; P < 0*.*0001)*. These results suggest that the starved L1 stage is optimal for cryopreservation of *C. elegans*, and that trehalose-DMSO solution is a more effective cryoprotectant than the freezing solution.

**Figure 1.**
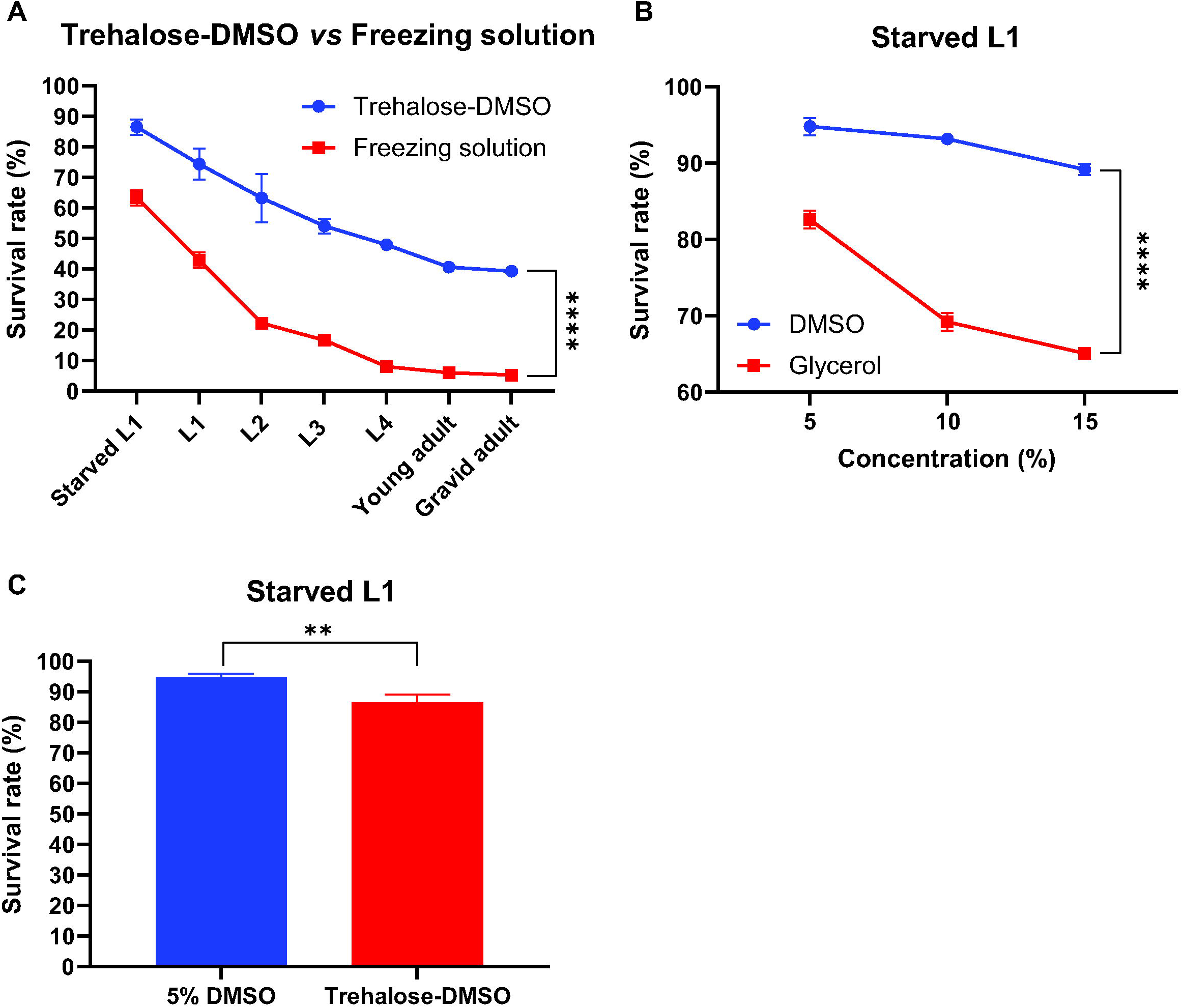
Systematic evaluation of developmental stage and cryoprotectant optimization for *C. elegans* cryopreservation. **(A)** Comparison of post-thaw survival rates of different developmental stages (starved L1, L1, L2, L3, L4, young adult, and gravid adult) of *C. elegans* cryopreserved in either trehalose-DMSO or freezing solution. Starved L1 larvae showed the highest survival rate under both cryoprotectant conditions, with trehalose-DMSO significantly outperforming the freezing solution across all stages. **(B)** Post-thaw survival of starved L1 larvae cryopreserved in different concentrations (5%, 10%, 15%) of DMSO or glycerol. Among these, 5% DMSO provided the highest survival rate. Data represented as mean□±□SEM from three independent experiments. Two-way ANOVA was applied to calculate statistical significance of survival curves. \*\*\*\**P*□<□0.0001 **(C)** Direct comparison of worm viability between 5% DMSO and Trehalose-DMSO, with 5% DMSO showing significantly higher cryoprotective efficacy. Data represented as mean ± SEM from three independent experiments. The two-tailed Student’s *t* test was applied to calculate statistical significance. \*\**P*□<□0.01.

### 3.2. Starved L1 larvae cryopreserved with 5% DMSO exhibit the highest survival rate

Glycerol and DMSO are the major cryoprotective agents used in standard freezing solution and trehalose-DMSO solution, respectively. We hypothesized that the concentration of the cryoprotective agents may influence the post-thaw survival of *C. elegans*, and by optimizing the cryoprotectant concentration, the cryopreservation protocol could be improved. To further optimize the concentration of cryoprotectants, we have cryopreserved the age-synchronized starved L1 larvae of wild-type N2 strain using different concentrations (5%, 10%, and 15%) of DMSO and glycerol. The samples were stored in a standard cardboard cryogenic box at -80 °C. After one week, the vials were thawed, and worm survival was scored. We observed that 5% DMSO resulted in the highest survival rate (∼95%), followed by 5% glycerol (∼83%), compared to higher concentrations of each cryoprotectant (Fig. 1B). Additionally, we observed that all concentrations of DMSO provided significantly better cryoprotection than the freezing solution *(F(1, 48) = 701*.*2; P < 0*.*0001)*. Moreover, the 5% DMSO significantly outperformed the trehalose-DMSO solution in promoting worm survival (Fig. 1C). These results indicate that 5% DMSO is the most effective cryoprotectant concentration for preserving starved L1 larvae of *C. elegans*.

### 3.3. Cryogenic storage container influences survival of starved L1 larvae during cryopreservation

To evaluate the effect of storage containers on cryopreservation efficiency, age-synchronized starved L1 larvae of the *C. elegans* N2 strain were cryopreserved in 5% DMSO using three different storage containers: a standard cardboard cryogenic box, a Styrofoam box, and an isopropanol-based freezing container. After one week of cryopreservation at –80□°C, the cryovials were thawed, and worm survival was scored. Among the three storage conditions, worms stored in the standard cardboard cryogenic box significantly exhibited the highest survival rate (∼91%, *P* < 0.0001) when compared to those stored in the Styrofoam box (∼31%) and isopropanol-based freezing container (∼6%) (Fig. 2A). These findings suggest that the type of storage container used during cryopreservation has a significant impact on worm viability. The standard cardboard cryogenic box provides optimal conditions for maintaining the survival of starved L1 larvae during freezing at -80 °C.

**Figure 2.**
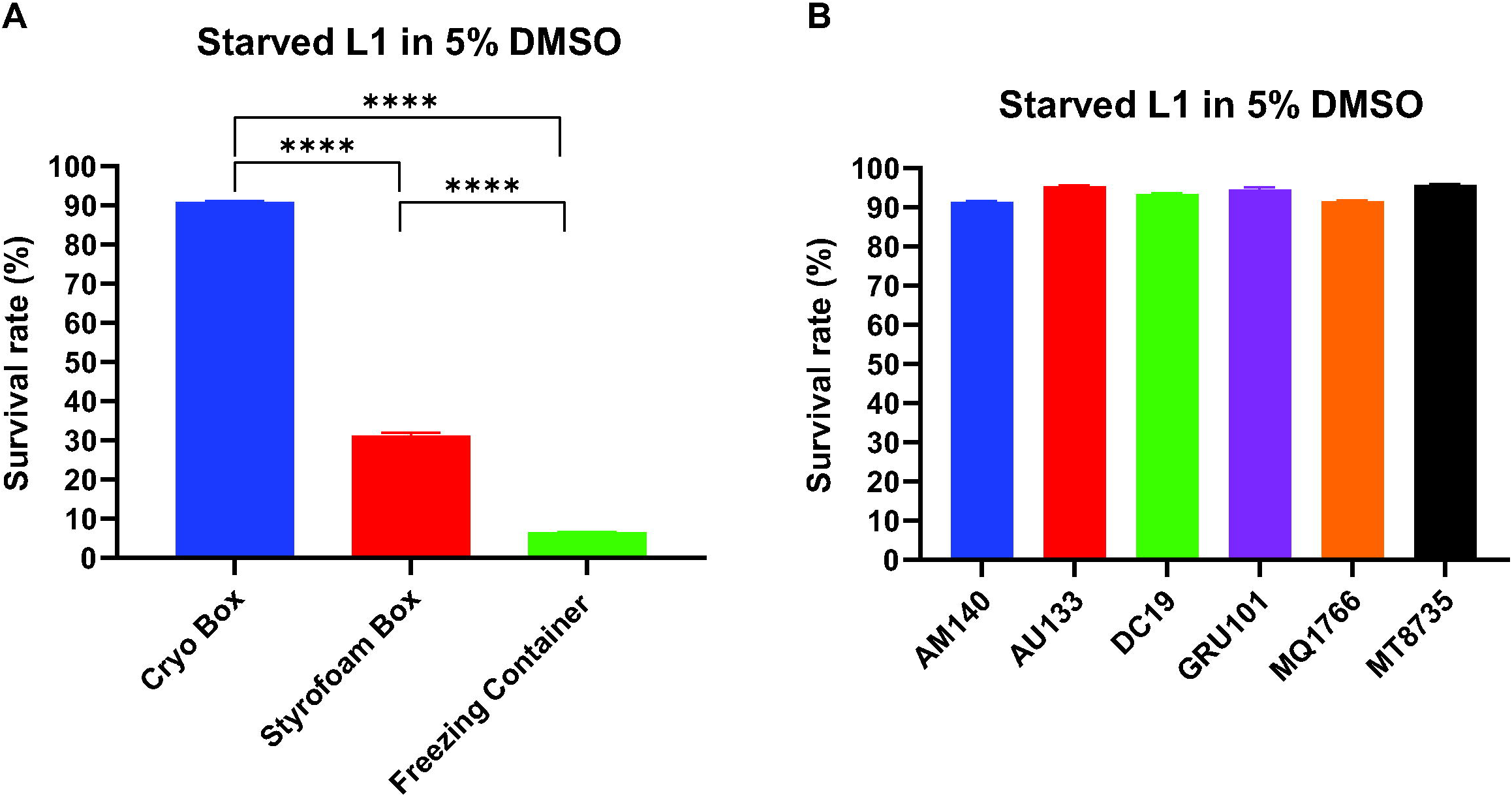
Influence of storage conditions and validation of optimized cryopreservation protocol across C. elegans strains. (A) Survival rate of starved L1 larvae cryopreserved with 5% DMSO stored in three different freezing containers: standard cardboard cryogenic box, Styrofoam box, and isopropanol-based freezing container. Worms stored in the standard cardboard cryogenic box showed significantly higher viability. Data represented as mean ± SEM from three independent experiments. The two-tailed Student’s *t* test was applied to calculate statistical significance. \*\*\*\**P*□<□0.0001. **(B)** Validation of the optimized cryopreservation protocol (5% DMSO, standard cardboard cryogenic box) across various mutant and transgenic *C. elegans* strains (AM140, AU133, DC19, GRU101, MQ1766, and MT8735). All strains consistently showed survival rates above 90%, confirming the broad applicability of the optimized protocol.

### 3.4. Mutant and transgenic C. elegans strains validate the optimized cryopreservation protocol

To validate the reproducibility and broad applicability of the optimized cryopreservation protocol using 5% DMSO, we assessed its effectiveness across multiple *C. elegans* mutant and transgenic strains available in our laboratory, including AM140, AU133, DC19, GRU101, MQ1766, and MT8735. Age-synchronized starved L1 larvae of each strain were cryopreserved in 5% DMSO using cryogenic vials stored in standard cardboard cryogenic boxes at –80□°C. After one week of storage, the vials were thawed at room-temperature, and worm survival was scored. We observed that all tested strains consistently exhibited a higher survival rate of > 90% (Fig. 2B). This suggests that the optimized cryopreservation protocol (Fig. 3) is robust, reproducible and broadly applicable to *C. elegans* strains with diverse genetic backgrounds.

**Figure 3.**
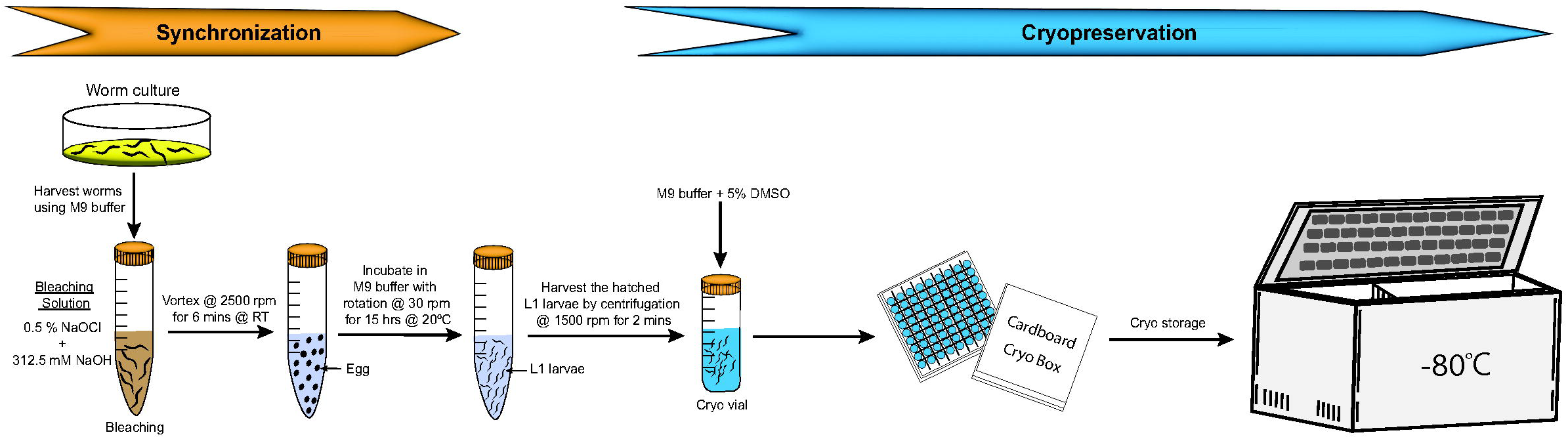
Workflow schematic and graphical summary of the optimized cryopreservation protocol. Schematic representation of the stepwise procedure for synchronized worm preparation and cryopreservation: starting from worm culture and bleaching for synchronization, hatching starved L1 larvae, followed by collection and cryopreservation in M9 buffer containing 5% DMSO. Cryovials are stored at - 80□°C in a standard cardboard cryogenic box.

## 3. Discussion

This study systematically evaluated the key factors influencing the cryopreservation efficiency of *C. elegans*, including development stage, cryoprotectant composition, and storage containers. Our results demonstrate that starved L1 larvae exhibited the highest post-thaw survival compared to fed larval stages (L1, L2, L3, L3, L4, young adult, and gravid adult) (Fig. 1A), corroborates earlier findings that starved worms and early larval stages survives better during cryopreservation^16,17^. Furthermore, we observed that the viability of the worm viability across all larval stages was significantly higher when cryopreserved using trehalose-DMSO compared to the conventional glycerol-based freezing-solution (Fig. 1A), consistent with the observations of Hayashi *et al*.^24^. Interestingly, trehalose-DMSO offered a better post-thaw survival rate in adult worms, including young adults (∼41%) and gravid adults (∼39%) when compared to the glycerol-based freezing solution (Fig. 1A), which has not been previously reported. These findings support to K. O’Connell’s proposition that trehalose-DMSO may offer superior cryoprotection over the glycerol-based freezing solution^25^. However, our results contrast with the observations of McClanahan *et al*., who reported that the post-thaw survival of early larval stages (L1-L3) was comparable to that of a mixed-stage populations, and that trehalose-DMSO and glycerol-based freezing solutions were similarly effective^22^.

Our results show that lower concentrations (5%) of both DMSO and glycerol provide greater post-thaw survival for starved L1 larvae of *C. elegans* compared to higher concentrations of 10% and 15% (Fig. 1B) Among these, 5% DMSO yielded the highest viability, outperforming 5% glycerol, an observation consistent with the previous findings of the Hayashi *et al*^24^. Moreover, when directly compared, 5% DMSO significantly outperformed the trehalose-DMSO mixture in preserving the viability of starved L1 larvae (Fig. 1C), which was not reported earlier, highlighting a potential refinement to existing protocols. Furthermore, our results demonstrate that cryovials containing starved L1 larvae in 5% DMSO placed within standard cardboard cryogenic storage boxes exhibited significantly higher post-thaw survival compared to those stored in Styrofoam boxes or isopropanol-based freezing containers (Fig. 2A). This observation contrasts with earlier recommendations, which suggest that slow freezing using Styrofoam or isopropanol-based containers enhances *C. elegans* viability during cryopreservation^17,21,22,24,25^. To our knowledge, no previous studies have systematically compared the influence of different storage containers on *C. elegans* cryopreservation, and we report this finding for the first time. Although the underlying cause of this discrepancy remains unclear, it may be attributed to differences in thermal conductivity, cooling rate consistency, or container-specific microenvironments that influence ice nucleation and cryoprotectant dynamics. These findings emphasize the need to revisit and systematically evaluate the physical parameters of storage containers, which are often overlooked yet may critically influence cryopreservation outcomes.

Most cryopreservation protocols for *C. elegans* have been developed and optimized using the wild-type N2 strain. However, there is a notable lack of studies reporting post-thaw survival outcomes for mutants or transgenic strains following these standard protocols. Exceptions are largely limited to investigations aiming to elucidate genetic mechanisms underlying freeze-thaw survival, such as studies involving insulin/IGF-1 signaling mutants like *daf-2* and *daf-16*^27^. As a result, researchers often lack insight into how well these cryopreservation protocols perform across genetically diverse or stress-sensitive strains, posing a risk of permanent loss of valuable mutants or transgenic lines. In this study, we addressed this critical gap by validating the optimized cryopreservation protocol that utilizes starved L1 larvae suspended in 5% DMSO and stored in cardboard cryogenic storage boxes. For the first time, this protocol was systematically evaluated across six different mutant and transgenic *C. elegans* strains, including stress-sensitive strains such as MQ1766 (sensitive to oxidative stress, osmotic stress, cold stress, and heat stress)^28^ and AM140 (sensitive to proteotoxic stress)^29^. Our results demonstrate that all tested strains exhibited high post-thaw survival rates exceeding 90% (Fig. 2B), highlighting the protocol’s broad applicability and robustness for preserving genetically modified *C. elegans* strains.

## 4. Conclusion

In this study, we systematically optimized a robust and reproducible cryopreservation protocol for *C. elegans*. Our results demonstrate that age-synchronized starved L1 larvae cryopreserved in 5% DMSO and stored in standard cardboard cryogenic boxes at -80□°C exhibit the highest post-thaw survival rates. This optimized condition consistently outperformed traditional freezing solutions. Furthermore, the protocol was validated across multiple mutant and transgenic strains, all of which exhibited high viability (> 90%), confirming its broad applicability and strain-independence. The simplicity, efficiency, and reliability of this protocol make it highly suitable for routine use in *C. elegans* research laboratories and genetic stock centers worldwide.

## Acknowledgements

The authors acknowledges the UPES, Dehradun, India, for providing institutional support, infrastructure and fundings for carrying out this research work.

## Author contributions

S.A. performed experiments and analyzed the results. A.K. performed experiments. K.R.B. designed and supervised the research work; wrote, edited and revised the manuscript.

## Declaration of interests

The authors declare no competing interests.

## Notes

### Competing Interest Statement

The authors have declared no competing interest.

